# Untangling the Components of Phenotypic Plasticity in Nonlinear Reaction Norms of *Drosophila Mediopunctata* Pigmentation

**DOI:** 10.1101/070599

**Authors:** Felipe Bastos Rocha, Louis Bernard Klaczko

## Abstract

Phenotypic plasticity may evolve as a generalist strategy to cope with environmental heterogeneity. Empirical studies, however, rarely find results confirming this prediction. This may be related to constraints imposed by the genetic architecture underlying plasticity variation. Three components of plasticity are central to characterize its variation: the *intensity* of response, the *direction* of response and the *total amount of change*. Reaction norm functions are a key analytical tool in plasticity studies. The more complex they are, the more plasticity components will vary independently, requiring more parameters to be described. Experimental studies are continuously collecting results showing that actual reaction norms are often nonlinear. This demands an analytical framework – yet to be developed – capable of straightforwardly untangling plasticity components. In *Drosophila mediopunctata*, the number of dark spots on the abdomen decreases as a response to increasing developmental temperatures. We have previously described a strong association between reaction norm curvature and across-environment mean values in homozygous strains. Here, we describe seven new reaction norms of heterozygous genotypes and further the investigation on the genetic architecture of this trait’s plasticity, testing three competing models from the literature – Overdominance, Epistasis and Pleiotropy. We use the curves of localized slopes of each reaction norm – Local Plasticity functions – to characterize the plastic response *intensity* and *direction*, and introduce a Global Plasticity parameter to quantify their *total amount of change*. Uncoupling plasticity components allowed us to discard the Overdominance model, weaken the Epistasis model and strengthen the support for the Pleiotropy model. Furthermore, this approach allows the elaboration of a coherent developmental model for the pigmentation of *D. mediopunctata* where genetic variation at one single feature explains the patterns of plasticity and overall expression of the trait. We claim that Global Plasticity and Local Plasticity may prove instrumental to the understanding of adaptive reaction norm evolution

## Introduction

Organisms may evolve the ability of producing different phenotypes in response to environmental variation – phenotypic plasticity – as a generalist adaptive strategy to cope with environmental heterogeneity [1]. Research on phenotypic plasticity has shown a remarkable increase in the last two decades [2]. Nonetheless, this growing effort has rarely detected unambiguous adaptive patterns of genetic variation of plasticity [3], even when the trait under study is clearly adaptive (e.g. [4, 5]).

In some cases, non-adaptive plasticity may even outnumber adaptive plasticity [6], suggesting that the evolution of an adaptive plastic response might be hindered by stringent constraints [7–10]. A possible source of constraint for the evolution of adaptive plasticity relates to the genetic architecture of the phenotypic response, which may be assessed by the study of reaction norms – the arrays of mean phenotypes produced by each genotype in response to a given environmental variable.

Scheiner [11] summarized three models concerning the genetic basis of phenotypic plasticity. The Overdominance model postulates that the amount of environmentally induced phenotypic change increases with the number of homozygous loci a genotype has. The Epistasis model explicitly assigns the control of the response to the environment to a set of loci that are independent from those controlling the expression of the trait across environments. The Pleiotropy model, in contrast, states that the response of a trait to the environment is controlled by the same gene(s) controlling the overall expression of the trait.

Three components of phenotypic plasticity are of central interest: *i*) the *intensity* of response (how fast does the phenotype change with the environment); *ii*) the *direction* of response (does it increase or decrease with the environmental variable); and *iii*) the *total amount of change* (the range of environmentally induced response). Characterizing these components may be more or less challenging, depending on the complexity of reaction norms.

If reaction norms are typically linear, the *intensity* and *direction* of response are constant. Moreover, the *total amount of change*, given a set of genotypes compared within the same environmental range, depends solely on the *intensity* of response: the higher the intensity of response of a genotype, the greater the total amount of change it will produce (Fig. 1A). If reaction norms, however, are nonlinear, the *intensity* of response becomes inconstant: a given genotype may be highly plastic at a given environmental range while being nearly non-plastic at another environmental range (Fig. 1B). Moreover, the *direction* of response may also become inconstant: a genotype may respond to increasing environmental values by producing higher phenotypic values and then at a given environmental value change its response producing lower phenotypic values (Fig. 1C). Therefore, the intensity and direction of response at a given environmental interval may not apply for another interval. Furthermore, the *total amount of change* is no longer directly deducible from the intensity of response at any environmental interval or even the mean intensity (Fig. 1D). It depends also on whether reaction norms are non-injective – i.e., have non-responsive segments (plateaus) and/or the *direction* of response is inconstant.

**Fig 1.**
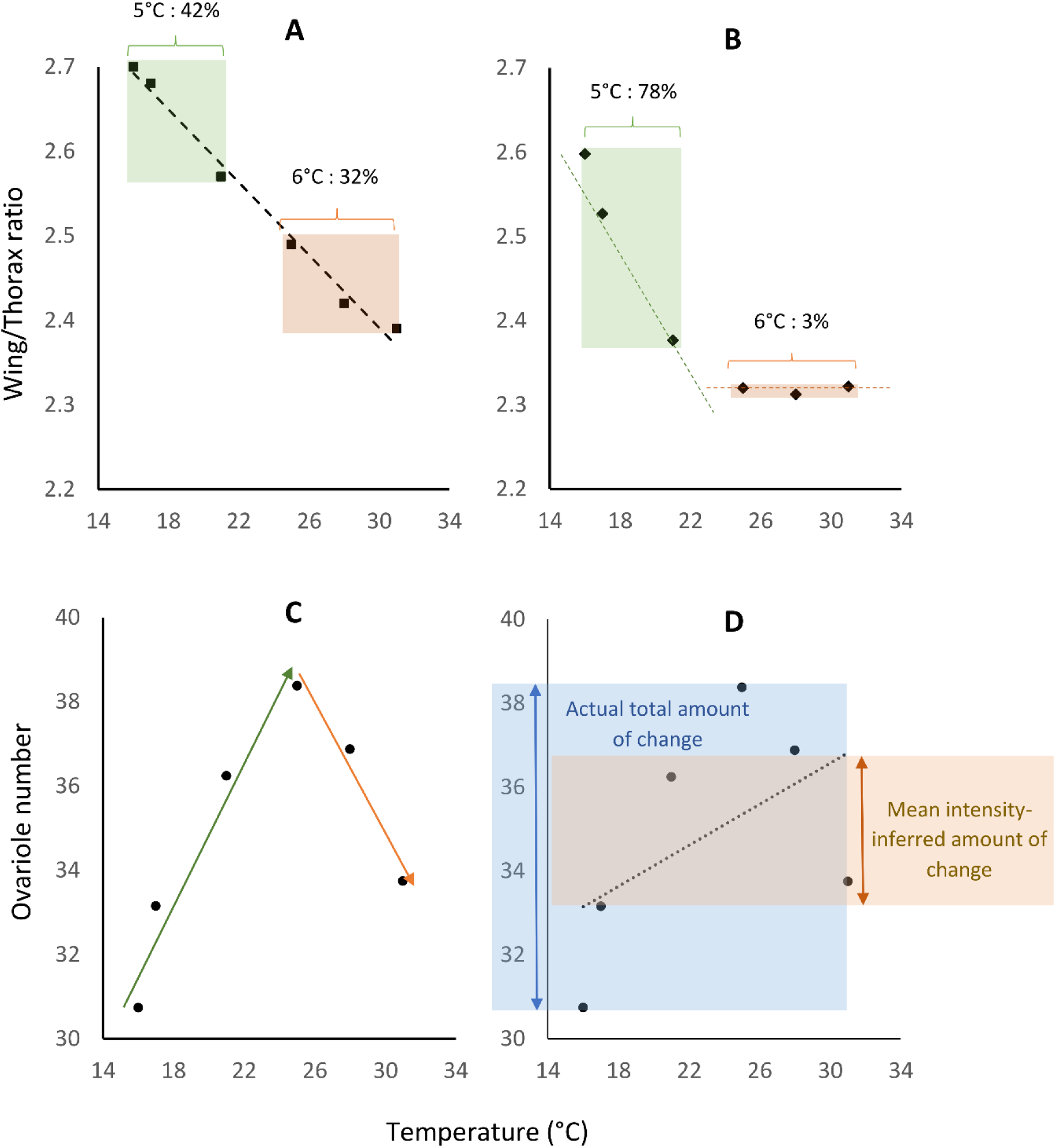
Thermal reaction norms of *Drosophila* (captured from Morin et al. 1997) illustrate the relationship between RN complexity and the components of phenotypic plasticity. A - Wing/thorax ratio reaction norm of *D. melanogaster*: the trait shows nearly constant intensity and direction of response across the whole thermal range, with approximately the same amount of change being observed within similar temperature intervals. The total amount of change may be reasonably estimated from either segments of the reaction norm. **B** - Wing/thorax ratio reaction norm of D. *ananassae*: the trait shows striking variation in response intensity, depending on the temperature interval considered. 78% of the total change is observed from 16 to 21°C, while from 25 to 31°C there is nearly no response. **C** - Ovariole number reaction norm of *D. ananassae*: the trait shows remarkable variation in the direction of response, increasing from 16 to 25°C and decreasing from 25 to 31°C (green and orange arrows, respectively). The linear model slope ignores the inconstancy of direction, describing the reaction norm as a continuous positive response to temperature. **D** - The non-injectivity of the ovariole number reaction norm breaks the direct relationship between the mean intensity of response (the mean slope) and the total amount of change: the total amount of chance inferred from the mean intensity of response is less than half the actual total amount of change.

Investigating the genetic architecture underlying phenotypic plasticity variation therefore requires taking two successive steps. The first one is determining what is the typical reaction norm shape. The second, given this typical shape, is determining how one may objectively characterize genetic variation at the relevant response features.

For simplicity, most plasticity studies model reaction norms as linear functions of the environment (i.e., P = g_0_ + g_1_E, where P is the mean phenotype, E is the environmental variable and g_0_ and g_1_ are genotype-specific coefficients) [e.g. 12–14]. Accordingly, experimental studies often quantify phenotypic plasticity by the difference between two phenotypes of a genotype submitted to two environmental conditions, or the slope of a linear equation adjusted to mean phenotypes from more environments [e.g. 15–17]. The central assumption of these studies, often implicit, is that a single parameter – the reaction norm slope – is sufficient to summarize all response features. If reaction norms are typically linear, such assumption does hold: the slope signal captures the *direction* of response; its absolute value gives the constant *intensity* of response (i.e., how much the phenotype is altered given one unit of environmental variable) and the product of the slope and the environmental range gives the *total amount of change*.

Since Krafka’s work [18], however, empirical studies describing reaction norms with more than two points along an environmental variable have provided mounting evidence that actual reaction norms are often nonlinear functions of the environment (e.g. [19–25]). Recently, studies using high-throughput RNA sequencing started to show that the same rule applies for gene expression at the whole-genome level. Chen et al. [26] found that 60.4% of the thermally responsive genes of *D. melanogaster* had quadratic reaction norms. Staton-Guedes et al. [25] observed that 75% of the thermally responsive genes of two ant species had a nonlinear response to temperature that “*would likely have been missed with a standard differential expression experiment (e.g. high vs. low temperature)*”.

In spite of the mounting evidence for the ubiquity of reaction norm nonlinearity and the biological relevance of reaction norm curvature, most plasticity studies persist on adopting a model whose central assumption is consistently falsified by empirical data. This persistent use of a clearly inadequate analytical framework may have led us to ignore biological meaningful dimensions of the plastic response, distorting the general picture we have of how phenotypic plasticity varies and evolves. Such distortion may be related to why, when theory predicts adaptive plasticity to be common, the body of empirical studies provides increasing evidence for its rarity.

The general nonlinearity of reaction norms poses the challenge of building an analytical framework capable of independently characterizing each component of phenotypic plasticity in an objective and straightforward manner. We have previously proposed the use of the localized slope (derivative) of a reaction norm – the Local Plasticity function of a reaction norm – to describe the trend of variation of response intensity and direction across the environmental axis [27]. A Local Plasticity function, however, does not quantify the total amount of change in a reaction norm, as reaction norms with different Local Plasticity functions may show the same overall response. A straightforward manner of quantifying this feature is taking the ratio between the phenotypic range – given by the difference between the maximum and minimum phenotypic values of each reaction norm – divided by the environmental range. Along with Local Plasticity functions, this parameter – Global Plasticity – may be used to characterize the relevant features of the plastic response described by the typically nonlinear reaction norm.

### Drosophila mediopunctata

Dobzhansky & Pavan, 1943 bears the characteristic pigmentation pattern of the *tripunctata* group: overall yellowish pigmentation with up to three distinct dark spots on the abdominal midline of the A4-A6 tergites [28]. The number of spots of *D. mediopunctata* is variable, ranging from zero to three. It is strongly affected by the second chromosome, where two inversions show contrasting patterns of association with the mean phenotype: *PA0* is associated with low number of spots and *PC0* with high number of spots [29].

Hatadani *et al*. [29] analyzed eighteen strains of *D. mediopunctata* whose second chromosome (carrying either the *PA0* or *PC0* inversions) had been extracted from wild-caught females and placed on a homogeneous genetic background. They observed strong temperature effect and karyotype-temperature interaction on the mean number of dark spots on the abdomen. Later on, Rocha *et al*. [21] used eight strains from this study [29] to investigate the genetic architecture of the thermal reaction norm shape and the mean trait value of this trait. They used a stratified sampling design to deliberately uncouple the mean trait value from the second chromosome inversions. Four strains were homozygous for *PA0* and four homozygous for the *PC0* inversion. Each group of homokaryotypic strains contained at least one strain belonging to each of two contrasting phenotypic groups: heavily-spotted (or dark) group (mean number of spots > 2.70); lightly-spotted (or light) group (mean number of spots < 1.62). The shape of the reaction norms could thus show: (*i*) no association with either the mean trait value or karyotype; (*ii*) mean trait value-independent association with the karyotype; or (*iii*) karyotype-independent association with the mean trait value.

Their results clearly fitted the third scenario: heavily-spotted strains had bowed upward reaction norms, lightly-spotted strains had bowed downward reaction norms, and this pattern was independent from the karyotype. Accordingly, the curvature of quadratic polynomials adjusted to each reaction norm was strongly correlated with the mean trait value [17], suggesting that reaction norm shape and mean trait value may be pleiotropically determined. These results seem to support the Pleiotropy model. However, none of the relevant response features (*intensity*, *direction* and *total amount of change*) is straightforwardly described by reaction norm curvature, although it has already been proposed to be useful as a plasticity parameter [30]. Furthermore, heterozygous genotypes could weaken the conclusions from [21]: they could show a reduced response, giving support for the Overdominance model; or yield a break in the mean trait value-curvature correlation – e.g. low mean trait values with bowed upwards reaction norms –, which could support the Epistasis model.

Here, we describe the thermal reaction norms of the number of dark spots on the abdomen of seven heterozygous genotypes of *D. mediopunctata.* We then include the eight reaction norms previously described [21] and estimate the Global Plasticity and Local Plasticity functions of the full dataset to further the investigation of the genetic basis of phenotypic plasticity of this trait. Our results show that the *overall amount of change* in each reaction norm varies as a nonlinear function of the mean trait value, and that the *intensity* of the plastic response varies both among and within reaction norms. The analysis of Global Plasticity rules out the Overdominance model, while the pattern of Local Plasticity variation weakens the Epistasis model and strengthens the Pleiotropy model.

We elaborate a developmental model for the genetic basis of the trait plasticity and overall expression, where genetic variation in one single property is sufficient to explain the variation of Global Plasticity, Local Plasticity functions and mean trait value. We discuss how the analysis of reaction norms and phenotypic plasticity may affect both the characterization of the relevant biological features and our understanding of how phenotypic plasticity varies and evolves. The acknowledgement of reaction norm nonlinearity appears as a first step to the adequate description of the genetic variation of plasticity and, perhaps, to the understanding of the phenomena underlying reaction norm shape and variation.

## Material and methods

### Description of thermal reaction norms

To enhance our dataset for additional tests of associations with the mean trait value, we characterized seven new thermal reaction norms of the number of dark abdominal spots of *D. mediopunctata*. This increased the heterogeneity of mean trait values and the total number of reaction norms (from 8 to 15).

### Crosses

We designed crosses to produce both homozygous and heterozygous genotypes for the second chromosome inversions *PA0* and *PC0*: *PA0*x*PA0* (GxI; IxH; GxH); *PC0*x*PC0* (OxX) and *PA0*x*PC0* (XxI; OxD; OxG), original strains in [29]. The average number of spots between each pair of parental strains ranged from 1.45 to 2.38. First instar larvae were collected from each cross and groups of 20 larvae were transferred to vials with 7 ml of growth medium. Vials were kept in eleven different temperatures between 14°C and 24°C, with 1°C increments, and two replicates for each cross. The thermal gradient apparatus is described in [22]. Flies were at least three days old by the time we counted the number of dark spots on the abdominal tergites A4 to A6.

### Statistical analysis

#### Curve fitting

The mean phenotype per temperature, i.e., the mean number of spots for each strain in each temperature, was first calculated by taking the mean value between replicates per sex, and thereafter by the mean between male and female mean phenotypes. A second order polynomial (P = *g_0_* + *g_1_* E + *g_2_* E^²^) was fitted to the set of eleven mean phenotypes per temperature of each reaction norm. Curve fitting was performed by least-squares nonlinear regression.

We tested whether the correlation between the curvature of quadratic polynomials adjusted to each reaction norm and mean trait value we had previously found held for two datasets: the seven crosses described here; and the pooled dataset including the eight previously described in [21] (amounting to fifteen reaction norms).

#### Mean trait value and phenotypic plasticity

We took the difference between the highest and the lowest mean phenotypes in each reaction norm (the reaction norm phenotypic range). We used the ratio between the phenotypic range divided by the environmental range (24-14°C) to quantify the genotype Global Plasticity and tested whether it was a function of the mean trait value.

Local Plasticity values for the set of ten intermediary temperatures (i.e., 14.5, 15.5, 16.5, …, 23.5°C) were estimated for the pooled dataset of fifteen reaction norms. Local Plasticity values were calculated both by empirical estimation, taking the stepwise differences between consecutive pairs of mean phenotypes [i.e., (P_15°C_-P_14°C_); (P_16°C_-P_15°C_); …; (P_24°C_-P_23°C_)] and by polynomial estimation, taking the first derivative of the quadratic polynomials fitted to each reaction norm (LP = *g_1_* + 2 *g_2_* T). We performed a multivariate regression to test whether each set of Local Plasticity values (empirical and polynomial estimates) could be described as a linear function of both the mean trait value (mtv) and temperature (*T*):

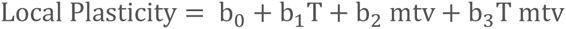

## Results

### Curvature and mean trait value variation

The crosses between the heavily-spotted and lightly-spotted strains previously described produced offspring with intermediary reaction norms (Fig. 2A and B). The quadratic model had a good fit for all reaction norms (R^²^ > 0.91), but for only two crosses it had a significantly higher fit than the linear model (GxI and IxH, Fig. 2B, solid lines) (p < 0.05, S1Table). For the other five the linear model had a sufficiently good fit (Fig. 2B dashed lines).

**Fig 2.**
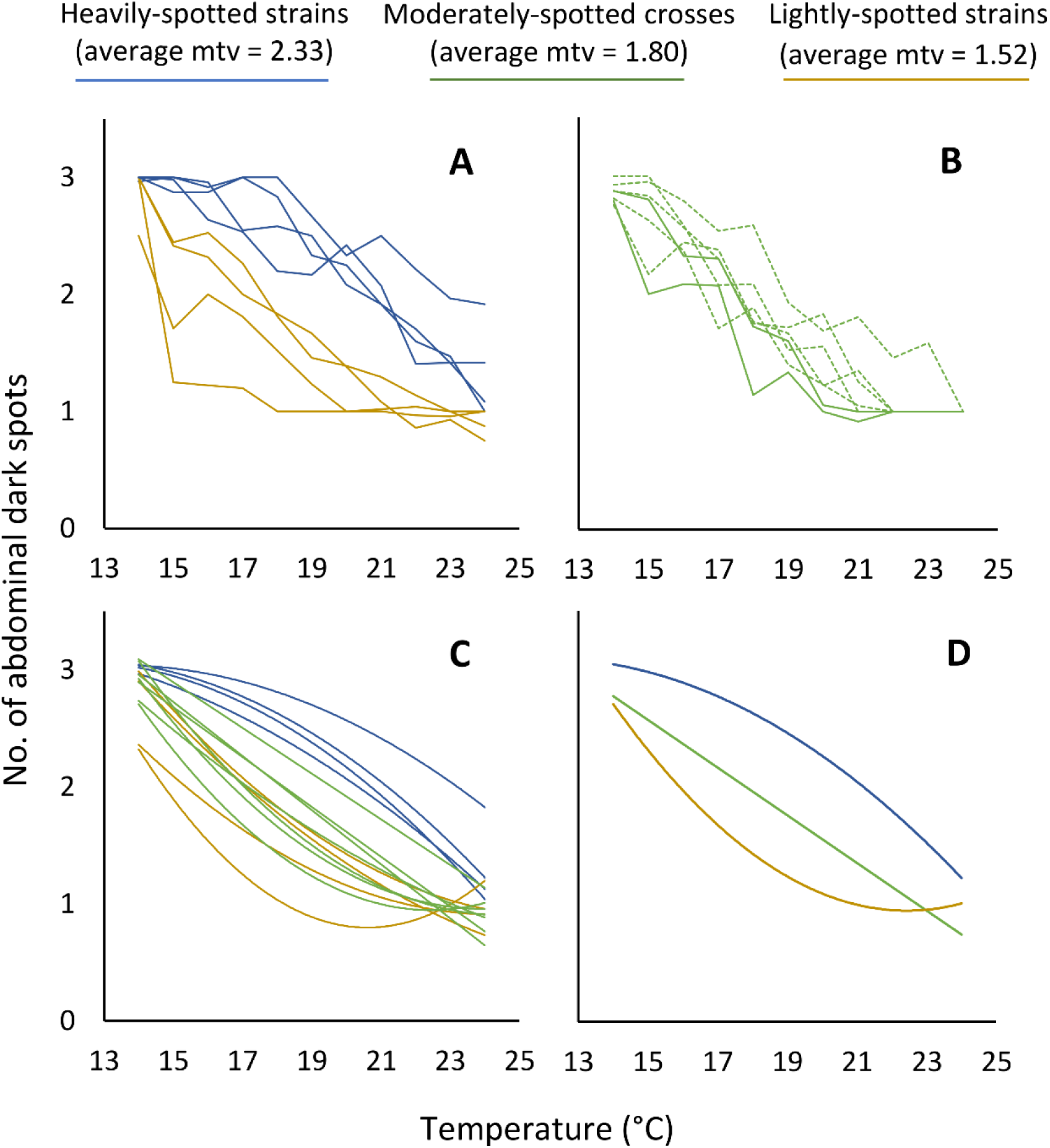
The shape of reaction norms of the number of abdominal spots of *D. mediopunctata* in response to temperature varies with the mean trait value. (A) Reaction norms of eight strains with contrasting mean trait value (mean trait value) and curvature described in [9] Rocha et al. (2009). (B) Reaction norms of the seven crosses performed to produce heterozygous genotypes (solid lines – reaction norms significantly best described by a second order polynomial). (C) Second order polynomials adjusted to each of the fifteen reaction norms. (D) Three reaction norms representative of the three groups of mean trait values tested: blue lines – heavily-spotted strains; green lines – moderately-spotted crosses; brown lines – lightly-spotted strains.

Nevertheless, the curvature of the newly described reaction norms was significantly correlated with the mean trait value (r = −0.92, p < 0.005) (Fig. 3A). When pooled with the eight reaction norms from [21] these values filled in the previous pattern of correlation, forming a continuous pattern where curvature is a linear function of the mean trait value. It had a negligible reduction in r value (from r = −0.93 to r = −0.92), but a very large increase in significance (from p < 0.001 to p < 0.00001) (Fig. 3B).

**Fig 3.**
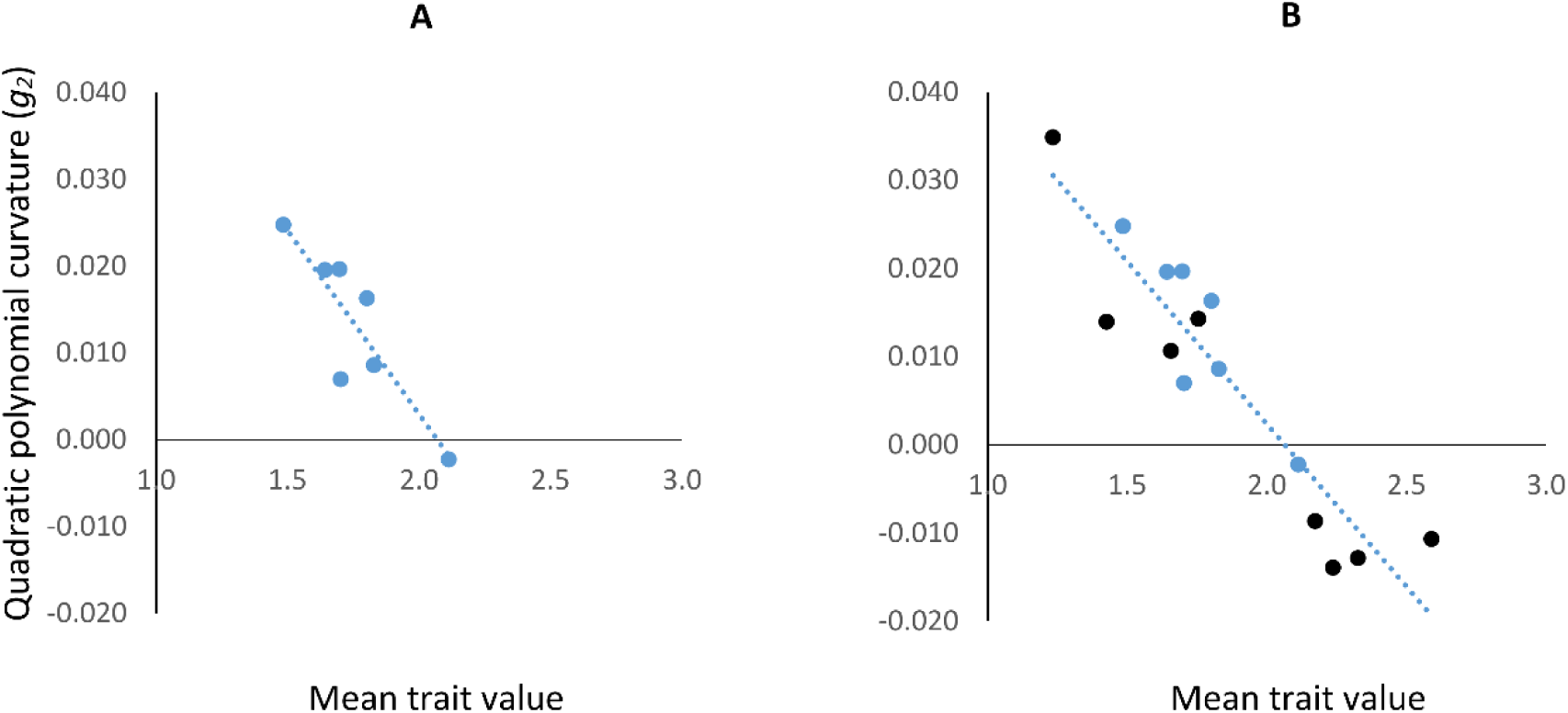
The curvature (*g^2^*) of second-order polynomials fitted to *D. mediopunctata* pigmentation thermal reaction norms is strongly correlated to the mean trait value. (A) Data only from the newly described crosses reaction norms; (B) data from the whole set of reaction norms including results from Rocha et al. (2009).

Therefore, the absolute curvature of a reaction norm may be interpreted as a measure of the inconstancy of phenotypic plasticity [27]. Accordingly, intermediary mean trait value reaction norms, i.e., those which were nearly linear, showed approximately the same intensity of response (constant plasticity) all over the temperature range, from low to high. Reaction norms with high mean trait value, which showed the lowest (most negative) curvature values, had a weak response to low temperatures (up to 17°C) and a strong response to high temperatures (above 21°C) (Fig. 4A). Reaction norms with low mean trait value, which had the highest (most positive) curvature values, showed the opposite pattern: a strong response to low temperatures and weak response to high temperatures (Fig. 4B). All reaction norms showed nearly the same intensity of response at intermediary temperatures (from 17 to 21°C) (Fig. 4A, B and C).

**Fig 4.**
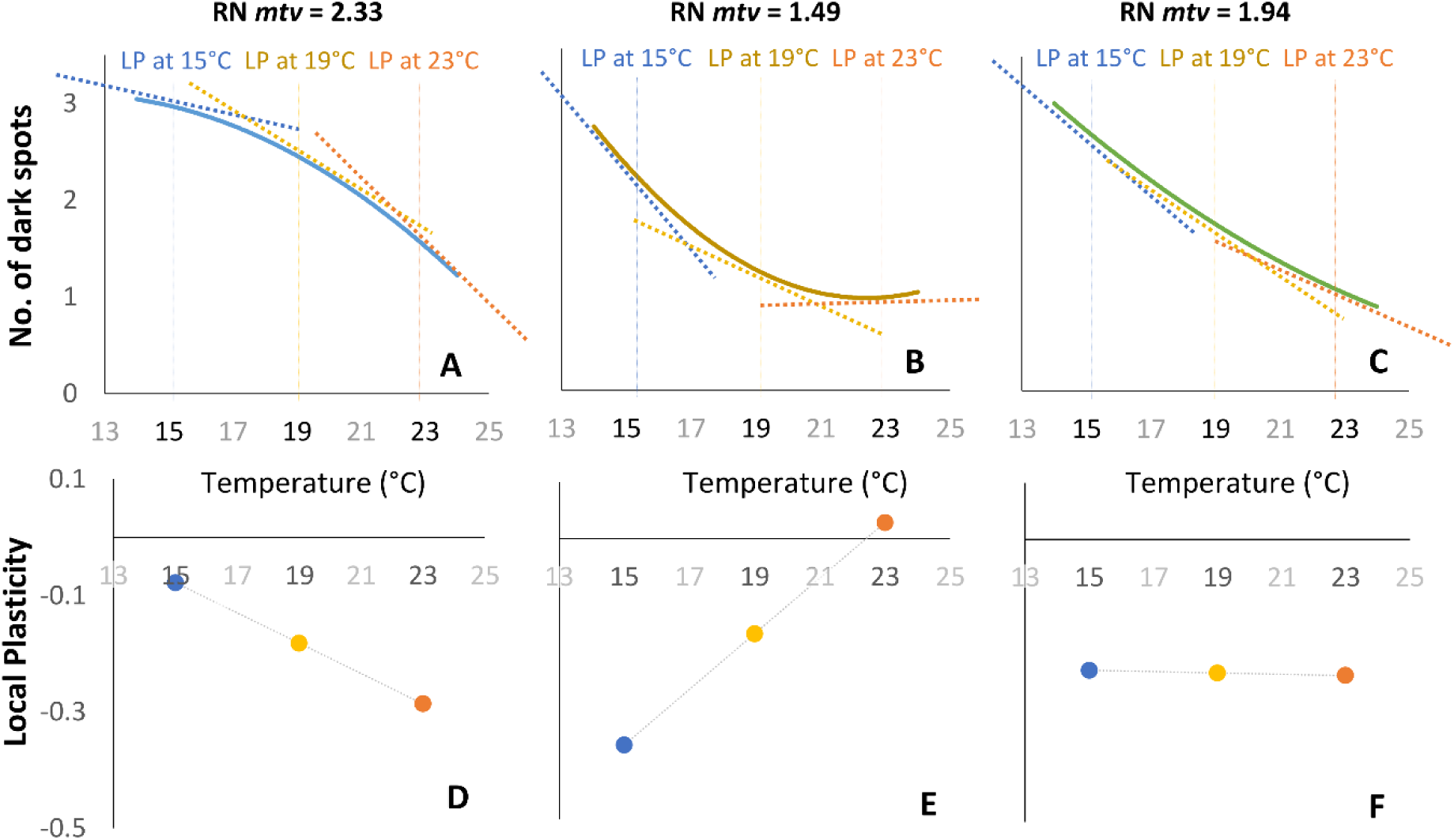
The tangent lines (whose slopes give the Local Plasticities) at three temperatures of reaction norms (RN) with contrasting mean trait values evidence the pattern of Local Plasticity variation as a function of the mean trait value (mtv). **A**, **B** and **C**: reaction norms of genotypes with high mean trait value (Z strain - blue line); low mean trait value (C cross – brown line); and intermediary mean trait value (G cross – green line). **D**, **E** and **F**: polynomial-estimated Local Plasticity functions of each reaction norm, highlighting the Local Plasticity values corresponding to the tangent lines shown above. Blue dotted lines and circles: 15°C. Yellow dotted lines and circles: 19°C. Orange dotted lines and circles: 23°C.

### Phenotypic plasticity parameters and mean trait value

Global Plasticity showed no evidence of linear variation with the mean trait value. Actually, it varied nonmonotonically with the mean trait value, being significantly described by a negatively curved parabola (p < 0.05, R² = 0.5329; Fig. 5): genotypes with more extreme mean trait values (predominantly homozygous) had lower Global Plasticity values and genotypes with intermediary mean trait values (predominantly heterozygous) had higher Global Plasticity values (Fig. 5).

**Fig 5.**
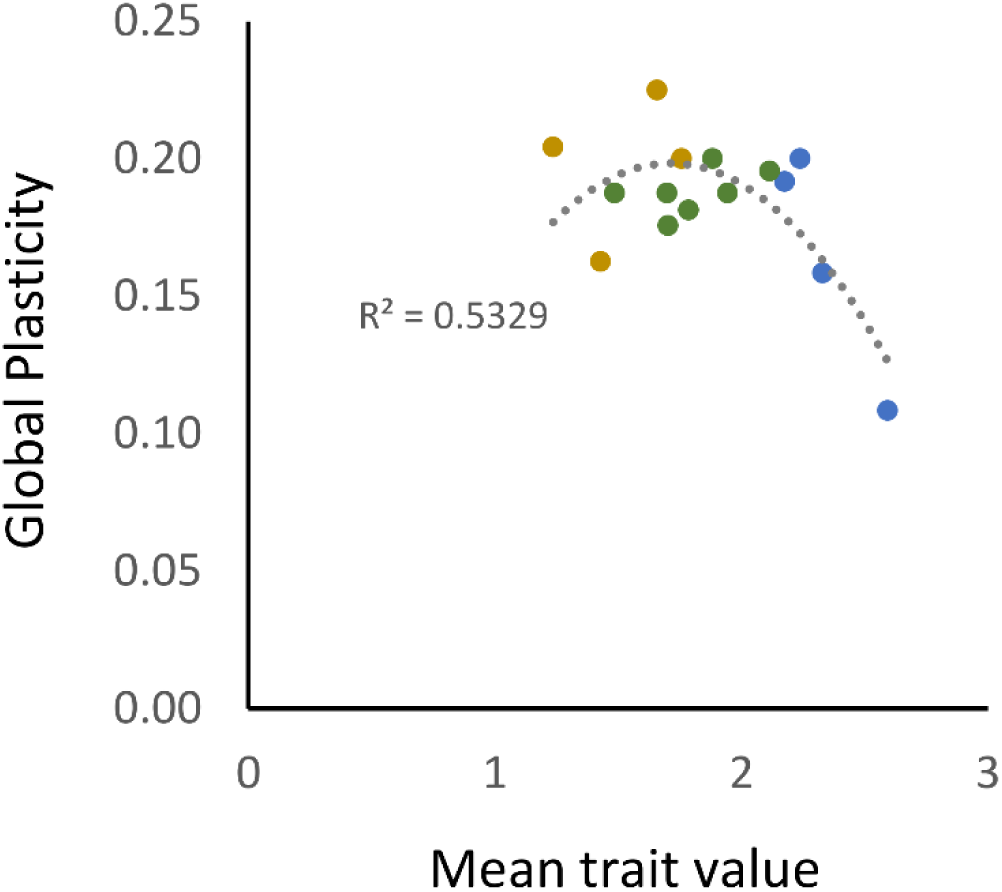
Global Plasticity values of fifteen reaction norms of *Drosophila mediopunctata* are a nonmonotonic function of the mean trait value (mtv). Green filled circles: heterozygous genotypes (described here); brown circles: homozygous strains with low mean trait value; blue circles homozygous strains with high mean trait value (data from Rocha et al. 2009).

The variation of Local Plasticity functions (i.e. the functions giving the localized response as a dependent variable of temperature) was tightly associated with the variation of mean trait value. Local Plasticity functions of genotypes with high mean trait value had negative slopes (Fig. 4D); Local Plasticity functions of genotypes with low mean trait value had positive slopes (Fig. 4E), while genotypes with intermediate mean trait value had nearly constant Local Plasticity values over all temperatures (Fig. 4F). The same overall result was obtained using either the empirical estimates or the polynomial-inferred Local Plasticity (see the general pattern given by the multivariate regression in S1 Figure and parameters in S2 Table).

### Discussion

The crosses described here were chosen to produce reaction norms of heterozygous genotypes, which showed intermediary mean trait value*s* and reaction norm shapes. When analyzed both isolated and pooled with the homozygous data, they clearly strengthened the pattern previously described, filling the gap between heavily-spotted and lightly-spotted reaction norms that supported the correlation between reaction norm shape and mean trait value. To our knowledge, this is the strongest demonstrated correlation of such type. It undoubtedly shows that the shape of these reaction norms should not be considered a separate trait from the pigmentation phenotype itself. This is in sharp contrast with the widely accepted [31] conjecture from Bradshaw that “*the plasticity of a character is an independent property of that character and is under its own specific genetic control*” [32].

### Analysis of phenotypic plasticity in nonlinear reaction norms

Our approach provides a means of breaking down the plastic response of nonlinear reaction norms into the key biological components of interest: Local Plasticity functions give the *intensity* and *direction* of response while Global Plasticity characterizes *the total amount of change*. Empirical estimation of Local Plasticity functions, however, demands experimentally challenging fine-scaled environmental gradients, as the one we used here. And estimation of Global Plasticity is highly dependent on environmental positioning if reaction norms are non-monotonic.

A possible alternative may be provided by polynomial reaction norm models [33] (P = g_0_ + g_1_E + g_2_E^2^ + … + g_m_E^m^ – where P is the phenotype of a trait, E is the environmental value and g_0_…g_m_ are genotype-specific coefficients). Previously, we proposed that quadratic polynomials could be “*a reasonable model for studies of reaction norm variation aiming to account for the nonlinearity of reaction norms*”, since they seem to provide the best balance between explanatory power and simplicity [22]. The Local Plasticity function of a quadratic reaction norm is a simple linear equation (LP = g_1_ + 2 g_2_ E) that captures its trend of response variation [27]. Our results show that, provided reasonable fit of the quadratic polynomial to the actual reaction norms, Local Plasticity functions estimated from adjusted quadratic polynomials yield equivalent results to the empirically estimated values (S2 Table, S1 Figure). Similarly, Global Plasticity calculation is possible from the quadratic polynomial coefficients and environmental maximum and minimum values (see. S1 Appendix).

Therefore, quadratic polynomials may provide a means of characterizing the variation of phenotypic plasticity in nonlinear reaction norms without requiring fine-scaled environmental gradients. Three-point-curve reaction norms are the minimum data set for quadratic polynomials to be fitted. Ideally, however, experimental studies should assess whether the polynomial model of choice is a good descriptor of the actual reaction norms. Hence, experimental reaction norm studies using the quadratic model should use at least four environmental values to test each genotype. Of course, if the quadratic polynomial has a poor adjustment to the reaction norms, Local Plasticity functions and Global Plasticity should be estimated by alternative means (e.g., empirically estimated or derived from more complex models).

### A pleiotropic model for the phenotypic plasticity of D. mediopunctata pigmentation

All reaction norms analyzed here were very well described by quadratic polynomials (the minimum R² was 0.9, with the exception of one strain described in [21]). In eight out of fifteen reaction norms the quadratic model showed a significantly higher fit than the linear. Moreover, although all reaction norms showed decreasing phenotypes as the temperature increased, six reaction norms were clearly non-injective, with plateaus at lower or higher temperatures. In terms of plastic response components, these reaction norms showed constant *direction* of response, significantly inconstant *intensity* of response within and among reaction norms, and had non-responsive segments (plateaus) that would bias the estimation of the *total amount of change* by the mean slope. The separate analysis of each plasticity component allowed us to directly test the predictions of each plasticity model for the trait studied.

The Overdominance model predicts that heterozygous genotypes would be better in “buffering” environmental perturbation, thus showing decreased overall response to the environment when compared to homozygous genotypes. The pattern of Global Plasticity variation does not conform to this expectation. Actually, the quadratic regression of Global Plasticity over mean trait value describes a complex pattern of variation where the total amount of thermally induced change increases towards moderately-spotted genotypes (mean trait value ∼ 1.75) and decreases towards extreme mean trait value values.

This pattern, however, does not seem to hold a simple relation with the second chromosome genotype. Both homozygous and heterozygous genotypes had reaction norms with Global Plasticity values on the higher half of the interval of observed Global Plasticity values (i.e., 0.166 < Global Plasticity < 0.225), while the only three Global Plasticity values on the lower half (0.108 < Global Plasticity < 0.166) were produced by homozygous genotypes (Fig. 5). The roots of the adjusted quadratic equation give the values of mean trait value where Global Plasticity would reach zero: mean trait value = 0.259 and mean trait value = 3.170. Therefore, the pattern described here predicts that only genotypes whose overall trait expression is either depleted (i.e., close to the minimum possible phenotype – zero) or saturated (i.e., close to the maximum possible phenotype – three) would be capable of producing a robust (or canalized) phenotype. The evidence we have so far thus suggests that if any pattern of association between Global Plasticity and heterozygosity exists, it is in opposition to the Overdominance model prediction.

All strains had their phenotype reduced as the developmental temperature increased (Fig. 2). Therefore, the *direction* of plastic response was the same for all genotypes across the whole thermal gradient, suggesting that temperature may play a general inhibitory effect on pigmentation on this species. This feature was characterized by the Local Plasticity functions yielding only negative Local Plasticity values within the environmental range tested (Fig. 4 and S1 Figure).

In contrast, the *intensity* of plastic response showed remarkable variation, both within and between reaction norms: the mean phenotype may be either strongly reduced or remain unchanged, given the same increase in temperature, depending on the reaction norm and on the temperature interval. This variation is clearly associated with the overall expression of the trait. Lightly-spotted genotypes are more plastic at low temperatures and less plastic at higher temperatures (Fig. 2, brown lines) and thus have positively curved (concave up) reaction norms. Heavily-spotted genotypes are less plastic at low temperatures and more plastic at high temperatures (Fig. 2, blue lines) and therefore have negatively curved (concave down) reaction norm. Moderately-spotted genotypes (now studied) have nearly constant plasticity across all temperatures (Fig. 2, green lines), yielding reaction norms with near zero curvature.

The significant multivariate regression of Local Plasticity by temperature and mean trait value shows that the slope of Local Plasticity functions (i.e., the curves giving Local Plasticity as a function of temperature) is different from zero and varies with mean trait value. Lightly-spotted genotypes have Local Plasticity functions with positive slope, approaching zero as the temperature increases (Fig. 4E). Heavily-spotted genotypes have Local Plasticity functions with negative slopes, diverging from zero as the temperature increases (e.g. Fig. 4D). Moderately-spotted genotypes have Local Plasticity functions with nearly zero slope: they are equally plastic across all temperatures (Fig. 4F). This tight association between plastic response *intensity* and mean trait value thus weakens the Epistasis model and provides solid straightforward evidence for a pleiotropic model for the overall expression of the trait and its response to temperature.

Our results may therefore be used to elaborate a developmental model that may explain the observed patterns, given a set of principles reasonably well supported:

1. **Temperature variation exerts a general and probably unavoidable effect on melanin synthesis:** lower temperatures promote melanization, while higher temperatures inhibit melanization. This principle is supported by experimental data from several *Drosophila* species [34–36], and seems to apply as well for *D. mediopunctata*, given the lack of variation in the *direction* of response.
2. **The upper limit for the number of dark abdominal spots in *D. mediopunctata* is three, and the lower limit is zero**. Even the most darkly pigmented flies observed so far were unable to produce dark spots on the A3 and A2 tergites (which would produce two additional phenotypes: four and five spots). Actually, this seems to be a phylogenetically inherited property, as the whole *tripunctata* group appears to show the same upper limit [28]. Obviously, no fly is able to have less than zero dark spots on their abdomen. Yet, males usually have at least one dark spot on A6, and thus the actual minimum may be slightly higher than zero.
3. **The lack of response in a given reaction norm is the result of either the saturation or depletion of the phenotype.** At temperatures where flies are induced to yield an overall level of melanization so high as to reach the upper phenotypic limit, reaction norms become unresponsive, yielding a plateau. At temperatures where melanization reaches so low levels as to reach the lower phenotypic limit reaction norms also yield a plateau, but at lower values.

If these principles are correct, the association between plastic response intensity (given by Local Plasticity functions) and the overall expression of the trait (given by mean trait value) may be explained by genetic variation at one single property: the overall production of melanin at the posterior tergites. Genotypes conferring high overall levels of melanin production (Fig. 2, blue lines) easily reach the maximum developmental limit (three spots) at colder temperatures, thus being less plastic. As the temperature increases, melanin synthesis inhibition reaches a point where spot formation becomes compromised, and these genotypes become plastic. Genotypes conferring low overall levels of melanin production (Fig. 2, brown lines) are only able to produce the maximum phenotype at very low temperatures, where melanin synthesis is highly promoted. A small increase in temperature, however, is sufficient to strongly inhibit their melanization, inhibiting spot formation and taking their reaction norms to the lower plateau, where these genotypes become less plastic. Genotypes conferring intermediary levels of melanin synthesis do not reach either limits and thus respond in a near linear fashion to the inhibitory effect of temperature. The portion of each reaction norm that forms a plateau (higher or lower temperatures), as well as the position of the plateau (at the upper or lower limit), would depend solely on the overall level of melanization conferred by each genotype interacting with underlying developmental limits and an unavoidable inhibitory effect of temperature on melanin synthesis.

### Global Plasticity, Local Plasticity and adaptive plasticity evolution

Our findings may have important consequences for the understanding of how an adaptive plastic response or, contrarily, an adaptive robust – or canalized – response may evolve. Global Plasticity and Local Plasticity functions may prove instrumental for future research on these questions, describing the relevant genetic variation in each case.

Adaptive phenotypic plasticity may be defined as the ability of organisms to produce specific phenotypic values as a response to specific environmental conditions. In the trait studied here, the variation of such fine-tuned response was not captured by Global Plasticity. Actually, reaction norm nonlinearity allowed genotypes to yield roughly the same total amount of change when raised across the thermal gradient while responding in remarkably contrasting manners. This contrast was clearly captured by Local Plasticity functions. For instance, they allowed us to distinguish between reaction norms that quickly drop from three to one spot between 14°C and 16°C (lightly-spotted genotypes) and those that remain nearly stable within the same segment (heavily-spotted genotypes). More importantly, they revealed that this property is strongly associated with the mean trait value.

Adaptive robustness may be described as the ability of genotypes to produce stable phenotypes in various environmental conditions. If the pleiotropic model outlined here applies, our results suggest that genotypes leading to either the saturation or depletion of the overall level of melanization would achieve a robust phenotype across temperatures. In contrast, genotypes conferring intermediary levels of overall melanization would produce an intense response to temperature (perhaps unavoidable, see [36]). This feature is not described by Local Plasticity functions, but is clearly revealed by the pattern of Global Plasticity variation.

It is worth noting that achieving higher levels of biological understanding on the genetic variation of phenotypic plasticity is a goal much more important than settling the discussion on a definite polynomial model for all reaction norms. Indeed, as argued in some instances (e.g. [37]), the slope of a linear model may be sufficient to characterize specific aspects of the plastic response. In the present case, it would suffice to characterize the constancy of response *direction.* Yet, it would fail to accurately describe the *total amount of change* in the non-injective reaction norms (i.e., those containing plateaus) and would simply erase the remarkable variation in plastic response *intensity*, falsely assigning a constant level of response to the genotypes with most extreme mean trait value values. The major harm of the use of a linear model in the present dataset, however, would be on the understanding of the biological system under scrutiny. Ignoring that reaction norms bend when reaching what appear to be the limits of the trait phenotypic space – which was captured by the quadratic polynomial curvature and Local Plasticity functions – we would not be able to elaborate the pleiotropic model outlined here, which may be used to further the investigation of this system.

## Acknowledgements

We thank Marcos R. D. Batista and Claudete Couto for technical help.

